# Genetic variation in the species *Arabidopsis thaliana* reveals the existence of natural heat resilience factors for meiosis

**DOI:** 10.1101/2024.07.16.603532

**Authors:** Jiayi Zhao, Huiqi Fu, Zhengze Wang, Min Zhang, Yaoqiong Liang, Xueying Cui, Wenjing Pan, Ziming Ren, Zhihua Wu, Yujie Zhang, Xin Gui, Li Huo, Xiaoning Lei, Chong Wang, Arp Schnittger, Wojciech P. Pawlowski, Bing Liu

**Affiliations:** Arameiosis Lab, South-Central Minzu University, Wuhan 430074, China; College of Life Science and Technology, Huazhong Agricultural University, Wuhan 430070, China; Department of Landscape Architecture, Zhejiang Sci-Tech University, Hangzhou 310018, China; College of Life Sciences, Zhejiang Normal University, Jinhua 321004, China; School of Public Health, Shanghai Jiao Tong University School of Medicine, Shanghai 200025, China; Shanghai Key Laboratory of Plant Molecular Sciences, Development Center of Plant Germplasm Resources, College of Life Sciences, Shanghai Normal University, Shanghai 200234, China; Department of Developmental Biology, University of Hamburg, Hamburg 22609, Germany; School of Integrative Plant Science, Cornell University, Ithaca, NY 14853, USA

## Abstract

Heat interferes with multiple meiotic processes leading to genome instability and sterility in flowering plants, including many crops. Despite its importance for food security, the mechanisms underlying heat tolerance of meiosis are poorly understood. In this study, we analyzed different meiotic processes in the Arabidopsis (*Arabidopsis thaliana*) accessions Columbia (Col) and Landsberg *erecta* (L*er*), their F1 hybrids and F2 offspring under heat stress (37°C). At 37°C, Col exhibits significantly reduced formation of double-stand breaks (DSBs) and completely abolished homolog pairing, synapsis and crossover (CO) formation. Strikingly, L*er* and L*er*/Col hybrids are much less affected than Col. Interestingly, only 10% ∼ 20% of F2 offspring exhibit the same heat tolerance of meiotic recombination as parents, indicating that heat resilience in L*er* is controlled by the interplay of several loci. Moreover, F2 offspring show defective chromosome condensation in interkinesis, and untimely sister-chromatid segregation and/or chromosome fragmentation, the levels of which exceed those in either inbreds and/or hybrids thus implying a transgressive effect on heat tolerance of meiosis. Furthermore, correlation and cytogenetic analysis suggest that homolog pairing and/or synapsis have an impact on heat tolerance of chromosome morphology and stability during post-recombination stages under heat stress. Taken together, this study reveals the existence of natural heat resilience factors for meiosis in Arabidopsis, which have the great potential to be exploited in breeding programs.

**Author summary:** Environmental temperature alterations affect meiotic recombination and/or chromosome segregation thus perturbing genetic makeup and genome stability in plants. We have previously reported that CO formation is fully abolished in *Arabidopsis thaliana* accession Col under heat stress (36°C-38°C) due to reduced DSB formation and impaired homolog pairing. Here, we show that in *Arabidopsis thaliana* accession L*er* under the same high temperature conditions, both DSB and CO formation occur normally, and homolog pairing is mildly impacted, which indicate a striking difference in heat tolerance of meiotic recombination from Col. Remarkably, Col/L*er* hybrids display the same heat tolerance as L*er*, however, only 10% ∼ 20% of F2 offspring behave the same as parents. Moreover, we found higher levels of defects in chromosome morphology and integrity, and sister-chromatid segregation in F2 population than those in both inbreds and hybrids, which suggest a transgressive effect influencing heat tolerance of meiosis. Our findings reveal that heat resilience in Arabidopsis is controlled by the interplay of multiple genomic loci, holding a great potential to be exploited in crop breeding.

## Introduction

Meiosis is a specialized type of cell division, which, in plants, generates spores with halved chromosome number required for sexual reproduction and maintenance of ploidy consistency over generations. During prophase I, homologous chromosomes (homologs) undergo recombination, resulting in reciprocal exchange of DNA fragments and thereby promoting genetic diversity. Meiotic recombination is initiated by the programmed formation of DNA double-strand breaks (DSBs) catalyzed by the conserved topoisomerase VI-like protein SPO11 together with other components of the DSB formation machinery (1, 2). The recombinases RAD51 and DMC1 catalyze DSB repair via homologous recombination (3–5). In plants, deficiencies in DSB repair result in impaired crossover (CO) formation and/or cause chromosome fragmentation (3–6). Although a large number of DSBs is generated along chromosomes, only a small proportion of them are repaired into COs. In Arabidopsis as in many other species, most COs are catalyzed by the ZMM proteins resulting in type-I COs that are placed at a distance from each other (CO interference) (7). The formation of the remaining approximately 15% of COs in plants involves structure-selective endonucleases such as MUS81 giving rise to type-II COs, which are typically not subject to CO interference (8). The process of COs formation can be visualized by the linking of homologs giving rise to bivalents connected by chiasmata. In the wild-type, all homologs become eventually connected through at least one CO (CO assurance) (9), which is required for equal distribution of homologs after meiosis I. During meiotic recombination, chromatids are organized into loops, which are anchored to the chromosome axis through the interactions between axis core proteins (e.g. ASY3 and ASY4 in Arabidopsis) and the axis-associated HORMA domain protein ASY1 (10–13). The assembly of the chromosome axis is required for normal DSB formation and repair, homolog pairing and synapsis, and recombination (12, 14–16).

The separation of homologs and sister chromatids is facilitated by spindles. Cohesin, a ring-shaped protein complex, assures the protection of the sister-chromatids union. The cohesion on chromosome arms is released during prophase and particularly during anaphase I allowing the separation of homologs (17). The centromeric cohesion is removed in anaphase II paving the road for the segregation of sister chromatids (18). The function and stability of centromeres are crucial for maintenance of cohesion and thus faithful chromosome segregation (19, 20).

Several studies have shown that meiosis in plants is sensitive to temperature variations (21–25). Either elevated or reduced temperature can interfere with cytoskeleton organization and/or meiotic cell cycle transition resulting in defective cytokinesis and unreduced spores (26–30). In Arabidopsis, mildly decreased or increased temperature within the spectrum that does not reduce fertility (8-28°C) promotes type-I CO formation (31, 32). Higher temperatures (30-34°C), however, attenuate homolog synapsis and alter CO rate and distribution, affecting meiosis progression (25, 30). In addition, increased temperatures perturb stability of centromeres and induce unbalanced chromosome segregation in Arabidopsis (33–35). At extremely high temperatures (36-38°C), homolog pairing and synapsis and CO formation in the Arabidopsis accession Columbia (Col) is completely abolished, which results in disrupted chromosome segregation and sterility (36–38). Despite the significance of heat stress on genetic makeup and genome stability in plants, the molecular mechanisms controlling heat tolerance of meiosis remain poorly understood.

Interestingly, previous studies have shown that there are naturally existing genetic differences with respect to meiotic recombination and stress tolerance in plants (39–45). Here, we report that the *Arabidopsis thaliana* accessions Col and Landsberg *erecta* (L*er*) show striking differences in heat tolerance of meiotic recombination. Our data reveal that meiotic chromosome stability in Col/L*er* F2 progeny is more sensitive to heat than in either inbreds or their hybrids indicating that several loci contribute to the natural heat resistance.

## Results

### Landsberg *erecta* exhibits higher heat tolerance of meiosis than Columbia

The *Arabidopsis thaliana* accessions Columbia (Col) and Landsberg *erecta* (L*er*), two frequently used varieties, exhibit differences in their meiotic recombination rates as well as response to stresses (39, 42, 45–48). To test whether heat tolerance of meiosis also varies, we compared the effect of extreme heat stress, i.e., 37°C, versus the control condition, i.e., 20°C, on these two accessions. To this end, we analyzed tetrad formation at the end of meiosis by orcein staining. At 20°C, both accessions generated typical tetrads that harbored four spores with equally-sized nuclei (Fig 1a and b). Exposure to the extreme temperature resulted in 85.7% tetrads in Col that showed irregular configurations with abnormal numbers and/or sizes of spores and nuclei (Fig 1a and b). In striking contrast, only 8.8% abnormal tetrads were found in L*er* (Fig 1a and b), suggesting that meiosis in L*er* is more resilient to heat stress than Col.

**Figure 1.**
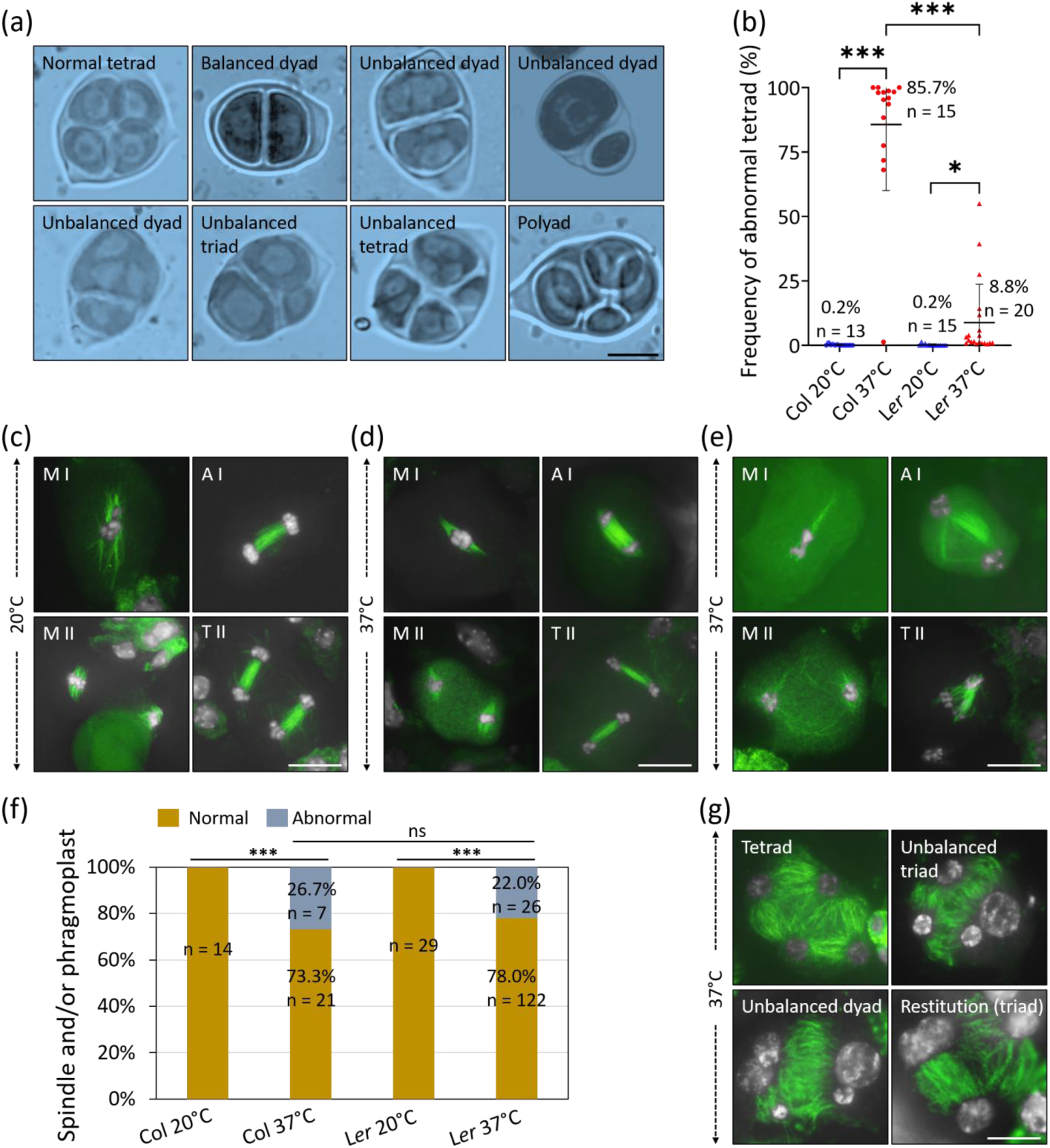
L*er* exhibits a higher heat tolerance of meiosis than Col. (a) Representative images of tetrads with normal or abnormal configurations. (b) Graph showing the frequencies of abnormal tetrads in Col and L*er* at 20°C or 37°C. Stars indicate significance levels based on unpaired *t* tests. (c and d) Representative images of immunolocalization of ɑ-tubulin in metaphase I, anaphase I, metaphase II and telophase II meiocytes depicting normal microtubule configurations at 20°C (c) or 37°C (d). (e) Representative images of immunolocalization of ɑ-tubulin in metaphase I, anaphase I, metaphase II and telophase II meiocytes displaying altered configurations at 37°C. (f) Graph showing the rates of spindle and phragmoplast with normal or abnormal configurations in Col and L*er* at 20°C or 37°C. Stars indicate significance levels based on χ^2^-tests. (g) Representative images of immunolocalization of ɑ-tubulin in tetrads showing normal or abnormal configurations at 37°C. White, DAPI; green, ɑ-tubulin. Frequencies represent the average rates of the corresponding phenotypes; n indicates the number of the analyzed plants or meiocytes; ***, *P* < 0.001; *, *P* < 0.05; ns, *P* > 0.05; scale bars, 10 μm.

Heat-induced formation of abnormal tetrads could result from defective meiotic recombination that leads to univalents and consequently, chromosome mis-segregation. Alternatively, defects in microtubule-mediated chromosome distribution and/or cytokinesis could be the underlying cause (38). To determine which meiotic process in Col is more affected by heat stress, we analyzed the spindle at metaphase I and II as well as phragmoplast at telophase II (27, 38, 49). At 20°C, the typical barrel-shaped spindles and straight phragmoplasts were observed in both accessions (Fig 1c). At 37°C, we did not observe obvious defects in microtubule organization in prophase I meiocytes in either accession (Fig S1) while Col and L*er* showed similar rates of irregularities in morphology of spindles and phragmoplasts in meiosis I and/or II (Fig 1d, normal; e, abnormal; and f) that likely contributed to an unequal segregation of chromosomes (Fig 1g). The higher heat tolerance of meiosis in L*er* versus Col is thus likely not caused by the difference in microtubule organization, but possibly occurs prior to metaphase I, i.e., meiotic recombination.

### DSB and CO formation in L*er* are largely resistant to heat

To assess the possible resilience of meiotic recombination towards heat in L*er*, we analyzed chromosomes by spreading and scored bivalent formation in Col and L*er* at 20°C and 37°C. At 20°C, five bivalents were present in diakinesis in both accessions (Fig 2a and c). While bivalent formation in Col was completely impaired at 37°C, this was not the case in L*er* (Fig 2a and c). To examine then the effect of heat on CO formation, immunolocalization of the type-I CO marker protein HEI10 was performed and the number of HEI10 foci on chromosomes in diakinesis was quantified. At 20°C, as previously reported, Col showed a higher number of HEI10 foci than L*er* with 11.0 foci versus 9.4, respectively (Fig 2b and d) (45). At 37°C, no HEI10 foci were detected in Col whereas L*er* showed an average of 10.5 HEI10 foci per meiocyte, being not significantly different from the number at 20°C (Fig 2b and d). Moreover, all analyzed anaphase I meiocytes in heat-stressed Col showed mis-segregation of homologs while only 41.5% unequal homolog segregation occurred in L*er* under heat stress (Fig 2e and f). Based on these data, we conclude that heat stress does not reduce CO formation in L*er*. Since heat stress has earlier been found to partially suppress CO formation by reducing DNA double-strand break (DSB) formation (36), we next analyzed the localization of the DSB repair recombinases RAD51 and DMC1, two widely used DSB markers (5, 50). Consistent with previous reports, both the numbers of RAD51 and DMC1 foci on zygotene chromosomes in Col were significantly reduced after heat treatment (Fig 2g-j) (36, 51). In L*er*, however, the focus numbers of both proteins were not significantly altered by heat (Fig 2g-j), suggesting that heat stress does not reduce DSB formation in L*er*.

**Figure 2.**
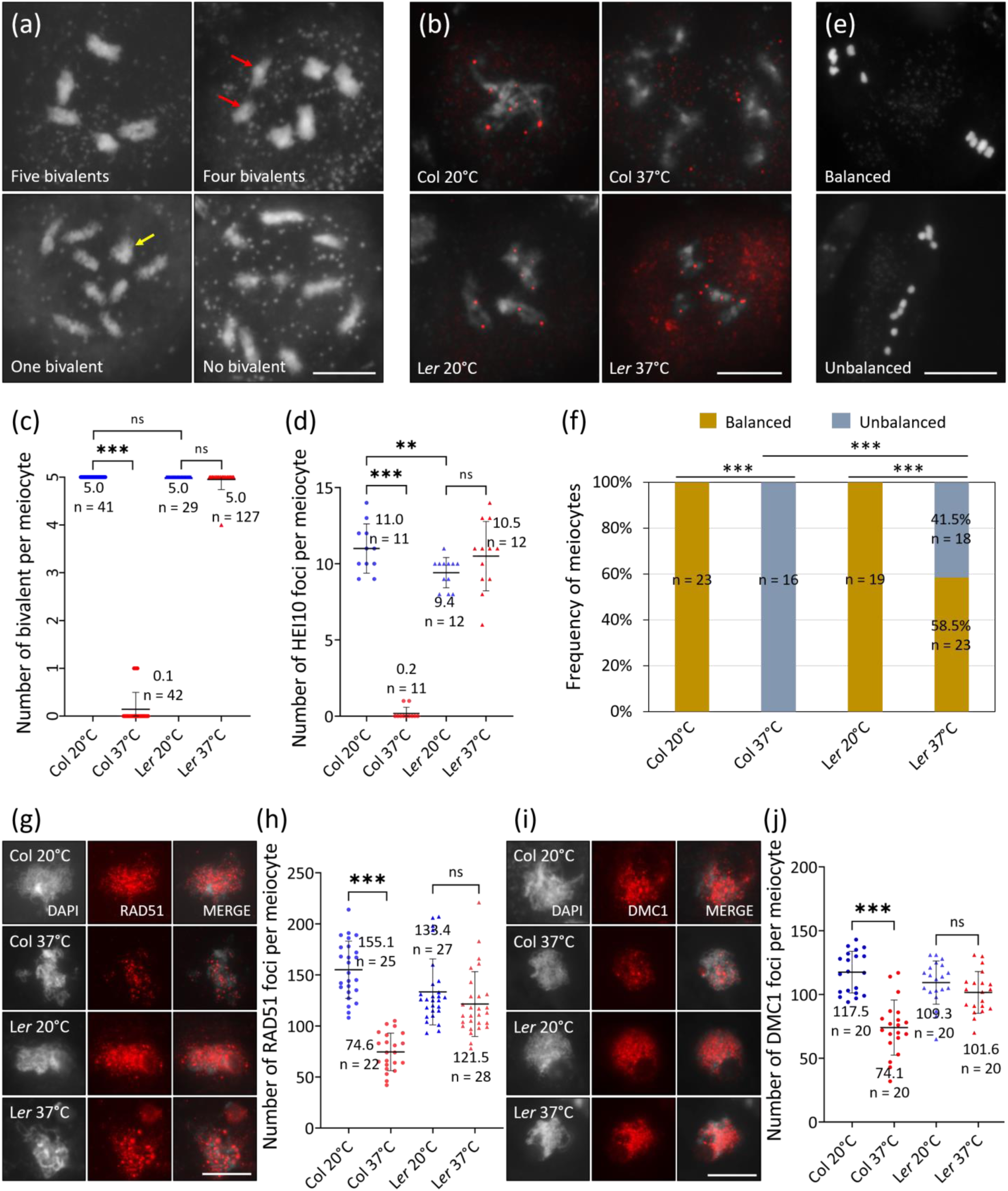
Heat stress does not reduce CO and DSB formation in L*er*. (a) Representative images of diakinesis meiocytes showing five, four, one or no bivalents. The red and yellow arrows indicate univalents and bivalent, respectively. (b) Immunolocalization of HEI10 in diakinesis meiocytes of Col and L*er* at 20°C and 37°C. (c and d) Graphs showing the average numbers of bivalent (c) and HEI10 foci (d) per meiocyte in Col and L*er* at 20°C and 37°C. Stars indicate significance levels based on unpaired *t* tests. (e) Representative images of anaphase I meiocytes showing balanced and unequal homolog separation. (f) Graph showing the fraction of anaphase I meiocytes with equal or unequal homolog separation in Col and L*er* at 20°C and 37°C. Stars provide significance levels based on χ^2^-tests. (g and i) Immunolocalization of RAD51 (g) and DMC1 (i) on zygotene chromosomes of Col and L*er* at 20°C and 37°C. (h and j) Graphs showing the average numbers of RAD51 (h) and DMC1 (j) foci per meiocyte in Col and L*er* at 20°C and 37°C. Stars indicate significance levels based on unpaired *t* tests; frequencies indicate the average number or fraction of the corresponding phenotypes; n indicates the number of the analyzed cells; ***, *P* < 0.001; **, *P* < 0.01; ns, *P* > 0.05; scale bars, 10 μm.

### Homolog synapsis in L*er* is more tolerant to heat stress than in Col

Disrupted homolog pairing is the main cause of impaired CO formation in Col under heat stress (36). Therefore, we compared heat tolerance of homolog pairing in Col and L*er*. We classified pachytene meiocytes into three types based on juxtaposition status of homologs: full-pairing, which indicates meiocytes exhibiting fully juxtaposed homologs (Fig 3a, b and e); partial-pairing, which indicates meiocytes showing both juxtaposed and unpaired regions between homologs (Fig 3f-i); and no pairing that represents impairment of homolog juxtaposition (Fig 3c and d). As previously reported, homolog pairing and synapsis in Col were fully impaired at 37°C (Fig 3a, c and j) (36). In L*er*, however, 63.1% of meiocytes exhibited full pairing (Fig 3e and j), and 35.2% of meiocytes showed partial pairing (Fig 3f-i, red arrows; j), with only 1.5% meiocytes displaying no pairing (Fig 3d and j). Next, we compared the impact of heat on chromosome morphology and the assembly of the chromosome axis and synaptonemal complex (SC) by analyzing the localization of ASY4 and ZYP1 on zygotene and pachytene chromosomes, respectively. At 20°C, ASY4 were loaded on the whole DAPI-stained chromosome regions showing a linear configuration (Fig 3k and l). Heat induced patchy ASY4 signals in both Col and L*er* (Fig 3k), but the fraction of this phenotype in Col was significantly higher than that in L*er* (Fig 3l, *P* < 0.001). Immunolocalization of ZYP1 showed that the SC assembly in Col was disrupted by heat, which, however, occurred normally or was only partially perturbed in most pachytene meiocytes in L*er* (Fig 3m-s, and t, ∼95.6%). These data indicate that homolog pairing and synapsis in L*er* are less impacted by heat than in Col.

**Figure 3.**
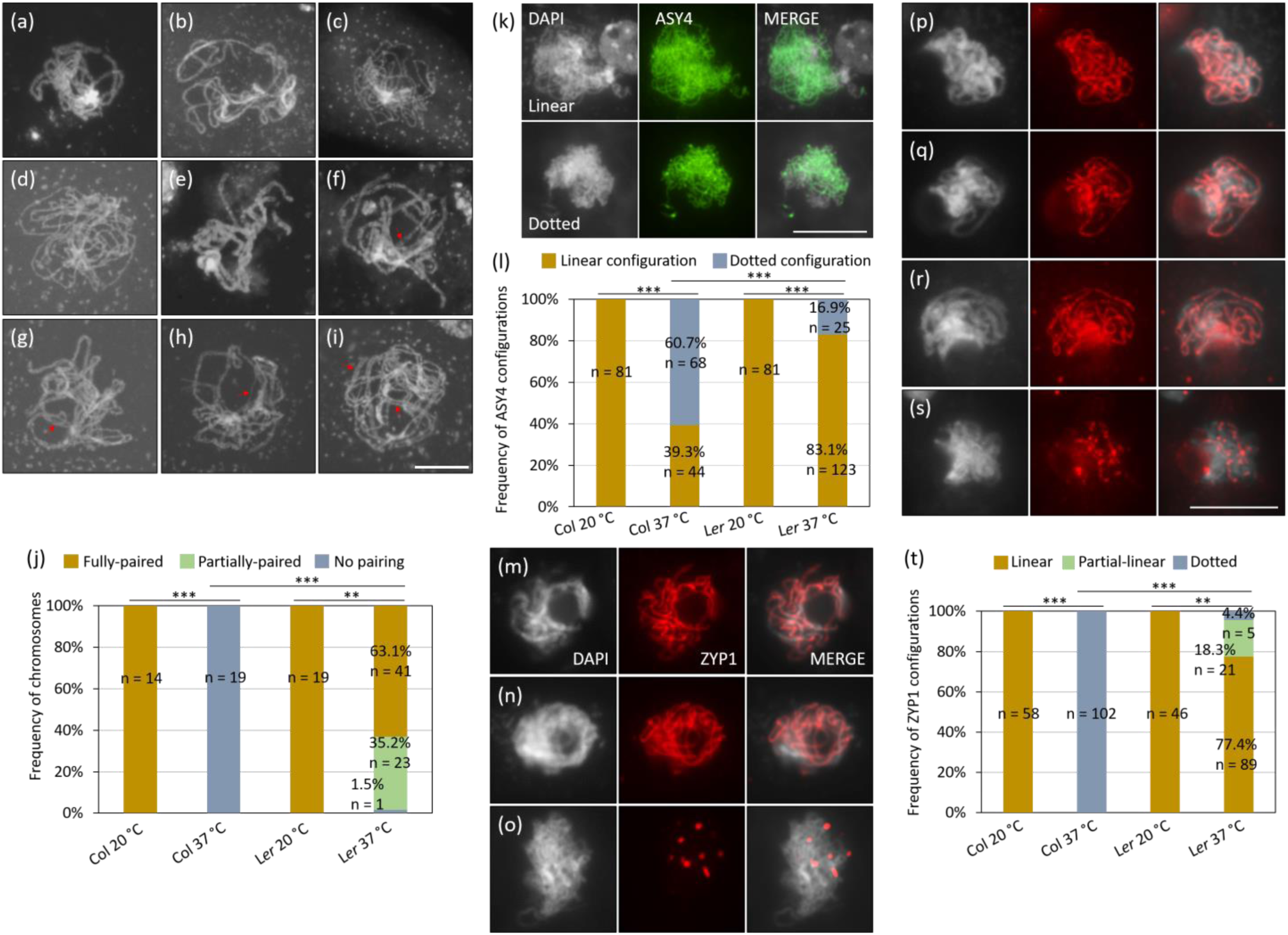
Homolog pairing and synapsis in L*er* are less impacted by heat than those in Col. (a-i) Pachytene chromosomes in Col (a and c) and L*er* (b, d-i) at 20°C (a and b) and 37°C (c-i). The red arrows indicate unpaired chromosome regions. (j) Graph showing the rates of pairing status of pachytene chromosomes in Col and L*er* at 20°C and 37°C. Stars indicate significance levels based on χ^2^-tests. (k) Representative images of immunolocalization of ASY4 on chromosomes showing linear or patchy configurations. White, DAPI; green, ASY4. (l) Graph showing the rates of ASY4 configurations in Col and L*er* at 20°C and 37°C. Stars indicate significance levels based on χ^2^-tests. (m-s) Immunolocalization of ZYP1 on pachytene chromosomes in Col (m and o) or L*er* (n, p-s) at 20°C (m and n) and 37°C (o-s). White, DAPI; red, ZYP1. (t) Graph showing the rates of ZYP1 configurations on pachytene chromosomes in Col and L*er* at 20°C and 37°C. Stars indicate significance levels based on χ^2^-tests; frequencies indicate the rates of the corresponding phenotypes; n indicates the number of the analyzed cells; ***, *P* < 0.001; **, *P* < 0.01; scale bars, 10 μm.

### Col/L*er* hybrids exhibit similar heat tolerance of meiotic recombination as L*er*

One of the main developmental differences between Col and L*er* is floral architecture (Fig S2), which, in Arabidopsis, is regulated by the receptor-like protein kinase ERECTA (ER) that acts upstream of the mitogen-activated protein kinase cascade module MKK4/5-MPK3/6 (52, 53). In addition, ER potentially plays a role in varied heat responses between Col and L*er* (54, 55). We thus examined whether the distinct heat tolerance of meiotic recombination between Col and L*er* is owing to a loss of *ER*. To this end, we analyzed bivalent formation, homolog separation, DMC1 and ASY4 localization, pachytene chromosome morphology as well as the SC assembly in Col plants defective for ER and/or MPK6. Both the single *er* and *mpk6* and the double *er mpk6* mutants displayed the same heat tolerance of meiotic recombination as Col (Fig 4d, f, h, j and n).

**Figure 4.**
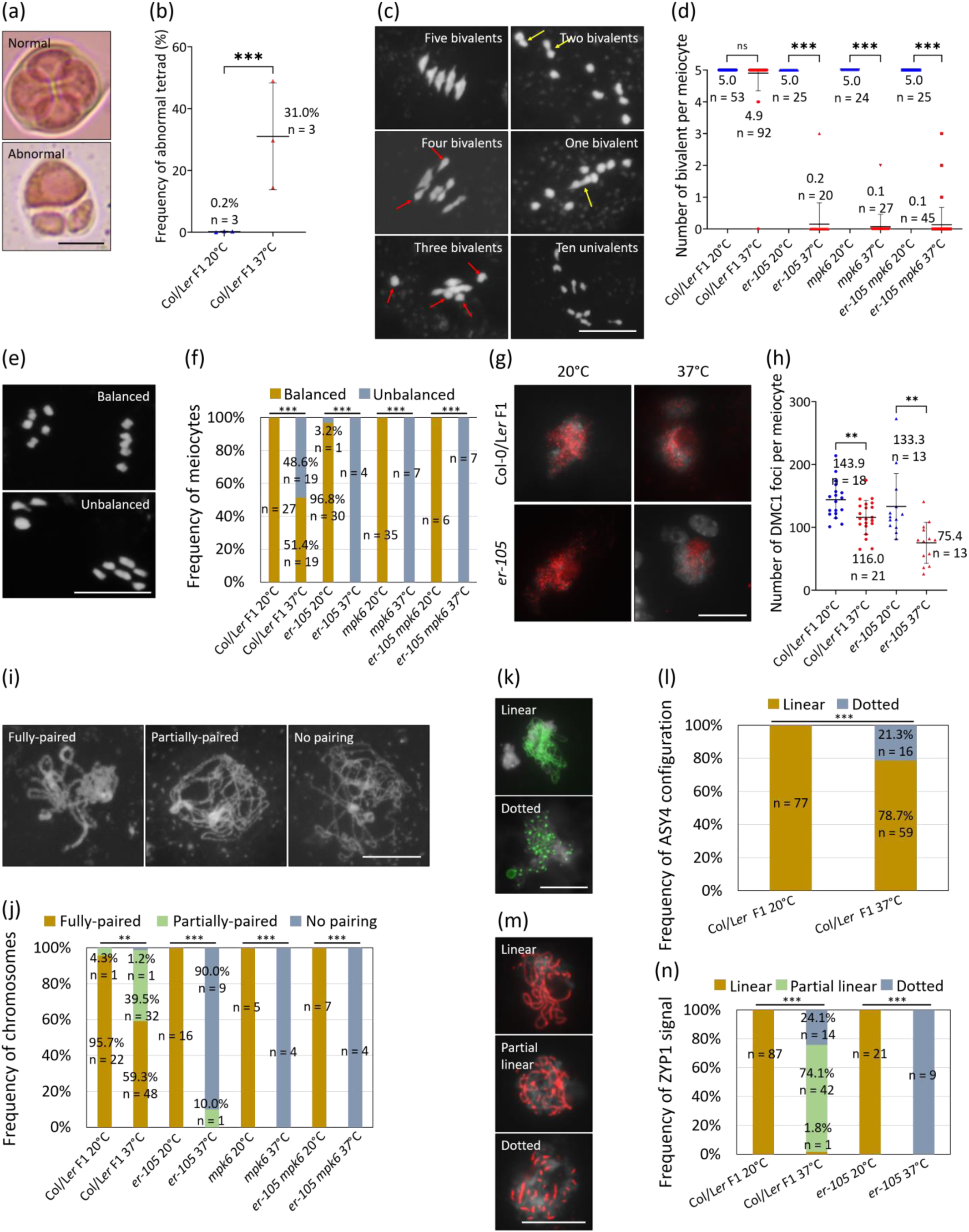
Col/L*er* hybrids exhibit similar heat tolerance of meiotic recombination as L*er*. (a) Representative images of normal and abnormal tetrads. (b) Graph showing the rates of abnormal tetrad in Col/L*er* hybrids at 20°C and 37°C. (c) Representative images of metaphase I chromosomes with different numbers of bivalents. The red and yellow arrows indicate univalents and bivalents, respectively. (d) Graph showing the average numbers of bivalent per diakinesis and/or metaphase I meiocyte in Col/L*er* hybrids, the single *er-105* and *mpk6*, and the double *er-105 mpk6* mutants at 20°C and 37°C. (e) Representative images of anaphase I chromosomes showing balanced or unbalanced homolog separation. (f) Graph showing the frequencies of anaphase I meiocytes with balanced or unbalanced homolog separation in Col/L*er* hybrids, the single *er-105* and *mpk6*, and the double *er-105 mpk6* mutants at 20°C and 37°C. (g) Immunolocalization of DMC1 on zygotene chromosomes in Col/L*er* hybrids and *er-105* at 20°C and 37°C. White, DAPI; red, DMC1. (h) Graph showing the average numbers of DMC1 foci per meiocyte in Col/L*er* hybrids and *er-105* at 20°C and 37°C. (i) Representative images of pachytene chromosomes showing full-pairing, partial-pairing, and no pairing. (j) Graph showing the rates of pachytene meiocytes exhibiting full-pairing, partial-pairing and no pairing in Col/L*er* hybrids, the single *er-105* and *mpk6* and the double *er-105 mpk6* mutants at 20°C and 37°C. (k) Representative images of immunolocalization of ASY4 on zygotene chromosomes showing different configurations. White, DAPI; green, ASY4. (l) Graph showing the frequencies of ASY4 configurations in Col/L*er* hybrids at 20°C and 37°C. (m) Representative images of immunolocalization of ZYP1 on pachytene chromosomes showing different configurations. White, DAPI; red, ZYP1. (n) Graph showing the rates of ZYP1 configurations on pachytene chromosomes in Col/L*er* hybrids and the single *er-105* mutant at 20°C and 37°C. In (b), (d) and (h), stars indicate significance levels based on unpaired *t* tests; in (f), (j), (l) and (n), stars indicate significance levels based on χ^2^-tests; ***, *P* < 0.001; **, *P* < 0.01; ns, *P* > 0.05; numbers or frequencies indicate the quantities or fractions of the corresponding phenotypes; n indicates the number of the analyzed individuals or cells. Scale bars, 10 μm.

To further explore the impact of the general genetic background on heat tolerance of meiosis, we analyzed chromosome behavior in heat-stressed Col/L*er* hybrids, which exhibited the same flower architecture as Col (Fig S2). These plants produced a lower level of abnormal tetrads (∼31.0%) than Col (Fig 4a and b; Fig 1b; *P* < 0.001). Interestingly, bivalent formation in Col/L*er* hybrids was not reduced under heat stress (Fig 4c and d), supporting the conclusion that heat tolerance of meiotic recombination in L*er* is not conferred by the flower architecture. In addition, balanced homolog segregation at anaphase I in heat-stressed Col/L*er* hybrids occurred at a similar frequency as in L*er* (Fig 4e and f; Fig 2f; *P* > 0.05). Moreover, at 37°C, zygotene chromosomes in Col/L*er* hybrids showed an average number of DMC1 foci (∼116.0) significantly higher than those observed for Col (∼74.1; *P* < 0.001) and L*er* (∼101.6; *P* < 0.05) (Fig 4g and h; Fig 2j). Analysis of pachytene chromosome configurations and ASY4 localization revealed no significant difference in heat tolerance between Col/L*er* hybrids and L*er* (Fig 4i-l; Fig 3j and l; *P* > 0.05). Furthermore, most pachytene meiocytes (∼74.1%) in Col/L*er* hybrids displayed a partial linear localization pattern of ZYP1, the extent of which was higher than that in L*er* (Fig 4m and n; Fig 3t; *P* < 0.001); and the meiocytes showing linear or patchy ZYP1 localization occupied 1.8% and 24.1%, respectively (Fig 4m and n). Overall, these findings revealed similar heat tolerance of meiotic recombination in Col/L*er* hybrids and in L*er*.

### Heat tolerance of pairing and CO formation is largely lost in the progeny of Col/L*er* hybrids

To further analyze the genetic underpinnings of heat tolerance of meiotic recombination, we examined homolog pairing, synapsis and bivalent formation in the progeny of Col/L*er* hybrids under heat stress. Unexpectedly, in a population of 1213 pachytene meiocytes from 132 Col/L*er* F2 individuals, only 11.1% of meiocytes showed full-pairing of homologs. Partial-pairing and no pairing of homologs occurred 36.7% and 52.2%, respectively (Fig 5a and b). Furthermore, only 19.8% of analyzed meiocytes (n = 3391) had five bivalents whereas 45.8% of meiocytes showed ten univalents, with the remaining 35% having either four, three, two or one bivalents occurring at roughly similar levels (Fig 5a and c). Accordingly, 48.4% of F2 individuals (n = 155) showed an average of at most one bivalent per meiocyte, while 18.7% of F2 individuals showed an average of more than four bivalents per meiocyte (Fig 5d). These data indicated that, in the context of bivalent formation, around half of the Col/L*er* F2 individuals behaved like Col, while only about 20.0% individuals displayed a L*er*-like resistance pattern under heat stress.

**Figure 5.**
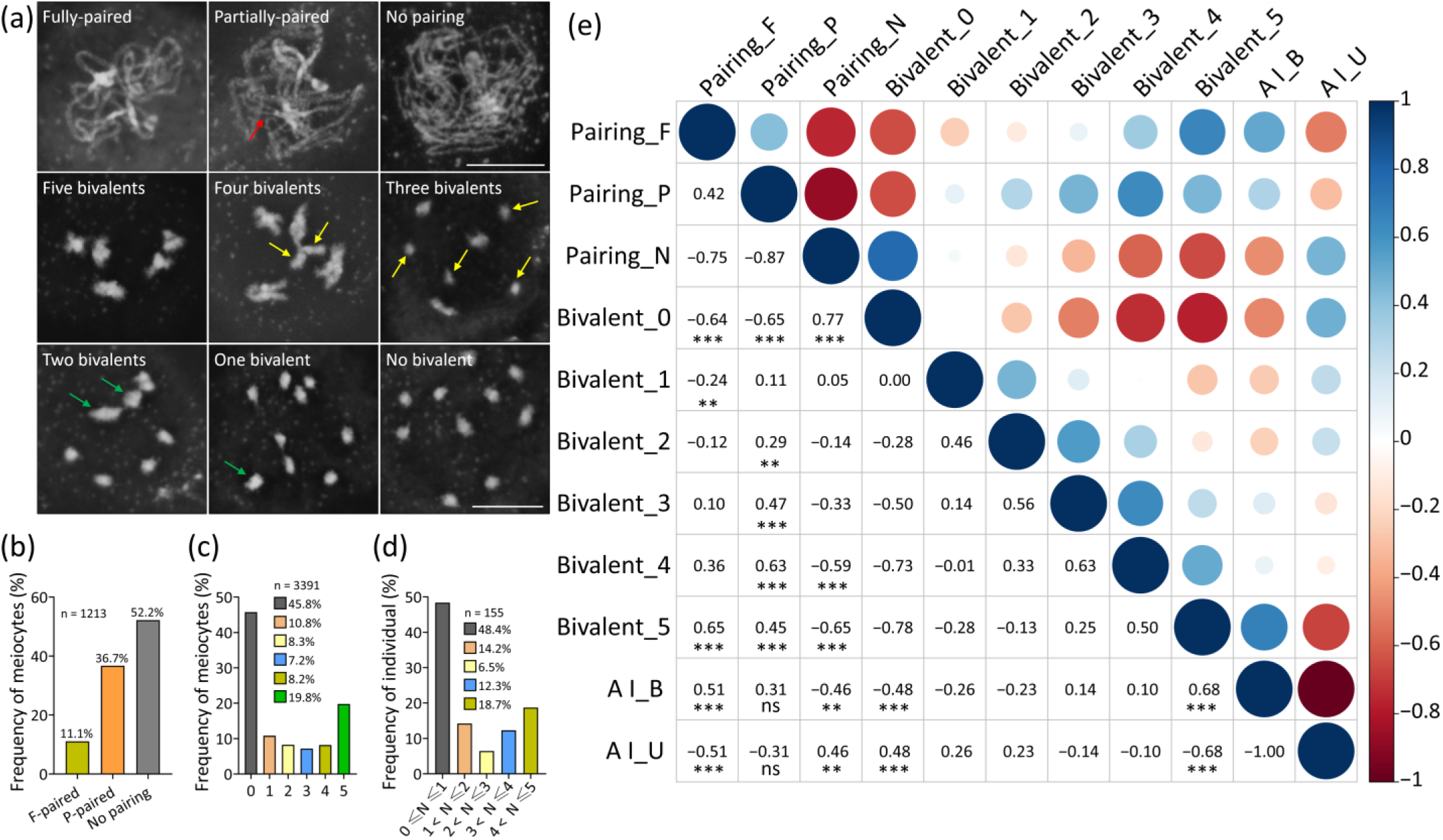
Chromosome pairing and bivalent formation in Col/L*er* F2 population under heat stress. (a) Representative images of chromosomes in pachytene and diakinesis showing different phenotypes of homolog pairing and bivalent formation in Col/L*er* F2 population under heat stress. The red, yellow and green arrows indicate unpaired or unsynapsed regions, univalents and bivalents, respectively. Scale bars, 10 μm. (b) Graph showing the fraction of pachytene meiocytes showing different pairing status in the heat-stressed Col/L*er* F2 population. (c) Graph showing the fraction of diakinesis meiocytes showing different numbers of bivalents in the heat-stressed Col/L*er* F2 population. (d) Graph showing the fraction of individuals in heat-stressed Col/L*er* F2 population showing different average numbers of bivalents. The frequencies indicate the rates of the corresponding phenotypes; n indicates the numbers of meiocytes or individuals; N indicates the average numbers of bivalent per meiocyte. (e) Graph showing correlation between pairing, average number of bivalents per meiocyte, and homolog segregation. Spearman correlation analysis was performed; numbers indicate the correlation coefficients between the corresponding phenotypes; significance levels of correlation were determined based on *t*-tests; ***, *P* < 0.001; **, *P* < 0.01; ns, *P* > 0.05; the significance levels in the cells without indication of significance values were not calculated.

We analyzed the correlation of the impact of heat on pairing, bivalent formation, and homolog segregation in the heat-stressed Col/L*er* F2 population by performing Spearman’s rank correlation tests using the frequencies of meiocytes showing different phenotypes (Fig 5e). As expected, a strong and positive correlation was found between the numbers of bivalents and homolog pairing (Fig 5e), with the presence of two, three or four bivalents per meiocyte being positively correlated with the partial-pairing phenotype (Fig 5e). This correlation offered a proof that our assay system worked. Moreover, positive correlations were found between balanced homolog segregation and bivalent formation or successful pairing (Fig 5e). Similar correlation patterns occurred in heat-stressed inbreds and F1 hybrids (Fig S3). These findings suggested that DSB formation and/or early stages of DSB repair-mediated homolog pairing are possibly the primary targets of heat.

### Heat stress reveals different impacts on chromosome morphology in interkinesis between Col and L*er* inbreds and their F2 offspring

At the end of meiosis I, homologs temporally decondense at interkinesis and then recondense at metaphase II (Fig S4). We found that heat induced a less-condensed or non-condensed chromosome configuration in interkinesis, which also varied between Col and L*er* (Fig 6a and c). The chromosomes exhibiting either kind of alterations in condensation showed a lower DAPI fluorescence intensity compared with the ones displaying normal condensation (Fig 6a and b). At 37°C, 25.0% Col plants produced meiocytes with a less-condensed chromosome configuration, while, this phenotype was not observed in any analyzed L*er* plant individuals (Fig 6c). Meanwhile, meiocytes showing a non-condensed chromosome configuration were observed in both Col and L*er* under heat stress (Fig 6c). Both kinds of abnormalities were observed in Col/L*er* hybrids and F2 population (Fig 6c). However, in the F2 offspring that displayed altered chromosome morphology, the fraction of meiocytes displaying non-condensed chromosomes was similar to that in Col but higher than those in L*er* and Col/L*er* hybrids (Fig 6d), while the fraction of meiocytes exhibiting the less-condensed phenotype was higher than those in inbreds and hybrids (Fig 6e).

**Figure 6.**
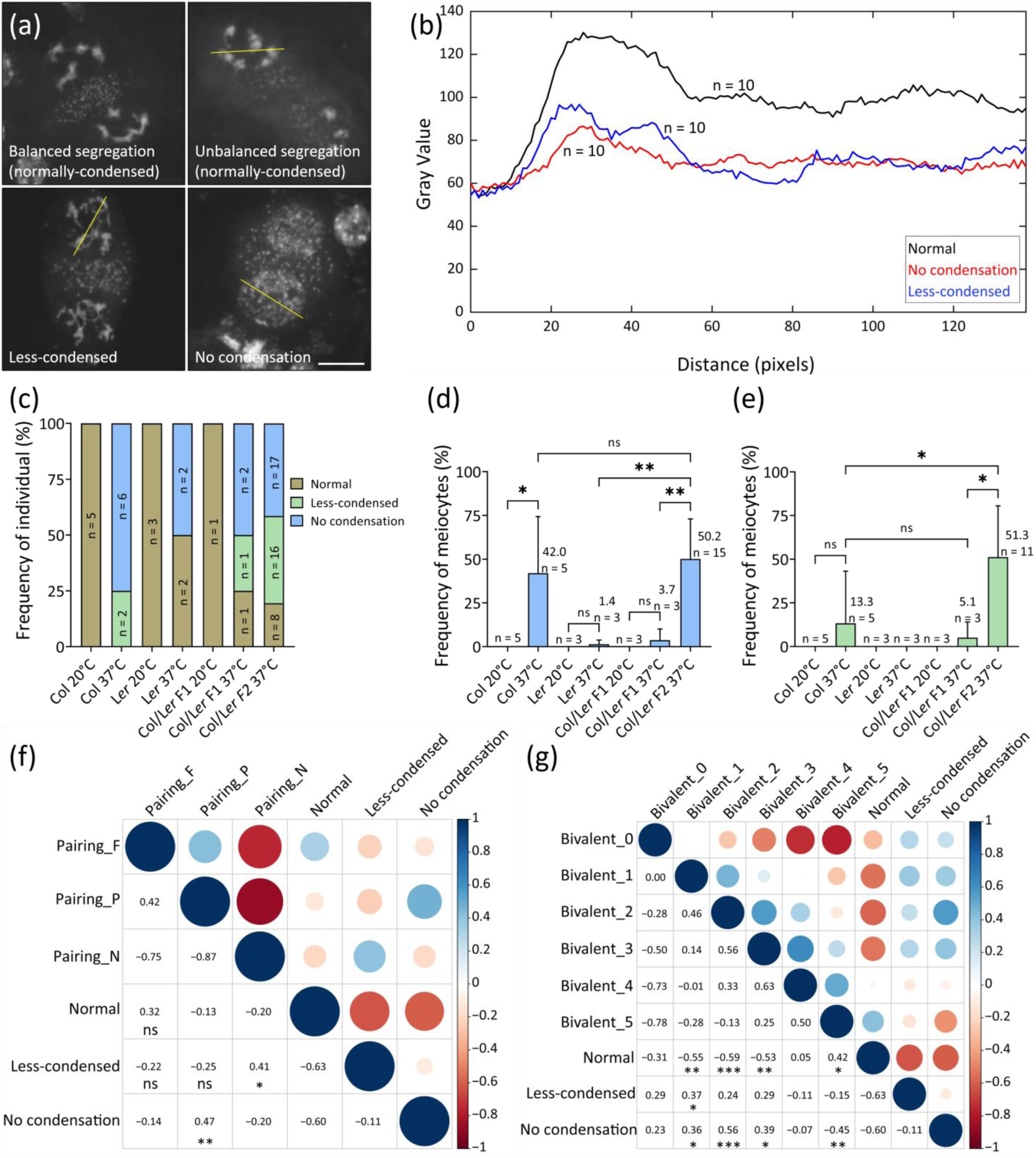
The impact of heat on chromosome morphology in interkinesis. (a) Representative images of different chromosome morphologies in interkinesis. Scale bar, 10 μm. (b) Average pixel intensity quantified in interkinesis meiocytes showing either kind of chromosome configuration of a section going through the middle of chromosomes (yellow lines). (c) Graph showing the rates of individuals with less-condensed or non-condensed phenotypes in Col, L*er*, Col/L*er* hybrids and Col/L*er* F2 at 20°C or 37°C. n indicates the numbers of plant individuals. (d and e) Graphs showing the rates of meiocytes showing non-condensed (d) or less-condensed (e) phenotypes in Col, L*er*, Col/L*er* hybrids and Col/L*er* F2 at 20°C or 37°C. Stars indicate significance levels based on unpaired *t* tests; n indicates the numbers of plant individuals; frequencies indicate the rates of the corresponding phenotypes. (f) and (g) Graphs showing the correlation between pairing (f) and average number of bivalents per meiocyte (g) with the less-condensed or non-condensed phenotypes. Spearman correlation analysis was performed; numbers indicate the correlation coefficients between the corresponding phenotypes; stars indicate significance levels based on unpaired *t* tests; ***, *P* < 0.001; **, *P* < 0.01; *, *P* < 0.05; ns, *P* > 0.05; the significance levels in the cells without indication of significance values were not calculated.

Interestingly, positive correlations were found between the less-condensed phenotype and impaired pairing or formation of only a single bivalent per meiocyte (Fig 6f and g). In addition, normal morphology of interkinesis chromosomes was highly correlated with the presence of five bivalents per meiocyte (Fig 6g). The non-condensed phenotype was positively associated with compromised pairing and reduced bivalent formation (Fig 6f and g). Taken together, these findings implied that the loci controlling heat tolerance of homolog pairing and/or CO formation may have a potential role in regulating chromosome morphology in interkinesis under heat stress.

### Heat stress induces high levels of chromosome instabilities in the Col/L*er* F2 population

Heat did not significantly increase the fraction of individuals exhibiting premature segregation of sister chromatids in Col and L*er* (Fig 7a-d). In the Col/L*er* F2 population, however, 56.7% of individuals (n = 164) showed ectopic segregation of sister chromatids from diakinesis to metaphase II (Fig 7b and c), suggesting that sister-chromatid cohesion was perturbed. In these defect-harboring F2 plants, the fraction of meiocytes showing untimely segregation of sister chromatids was significantly higher than in parental genotypes (Fig 7d). While we did not observe obvious defects in chromosome integrity in Col and L*er* under heat stress, 27.0% of individuals (n = 126) in the heat-stressed Col/L*er* F2 population produced chromosome fragments at metaphase I, anaphase I or metaphase II (Fig 7e-g), suggesting that DSB repair was defective in those F2 individuals under heat stress. Moreover, the average number of fragments per meiocyte in those F2 individuals was higher than in the inbreds and hybrids (Fig 7g). These data suggested that heat tolerance of chromosome stability maintenance is affected in the Col/L*er* F2 population.

**Figure 7.**
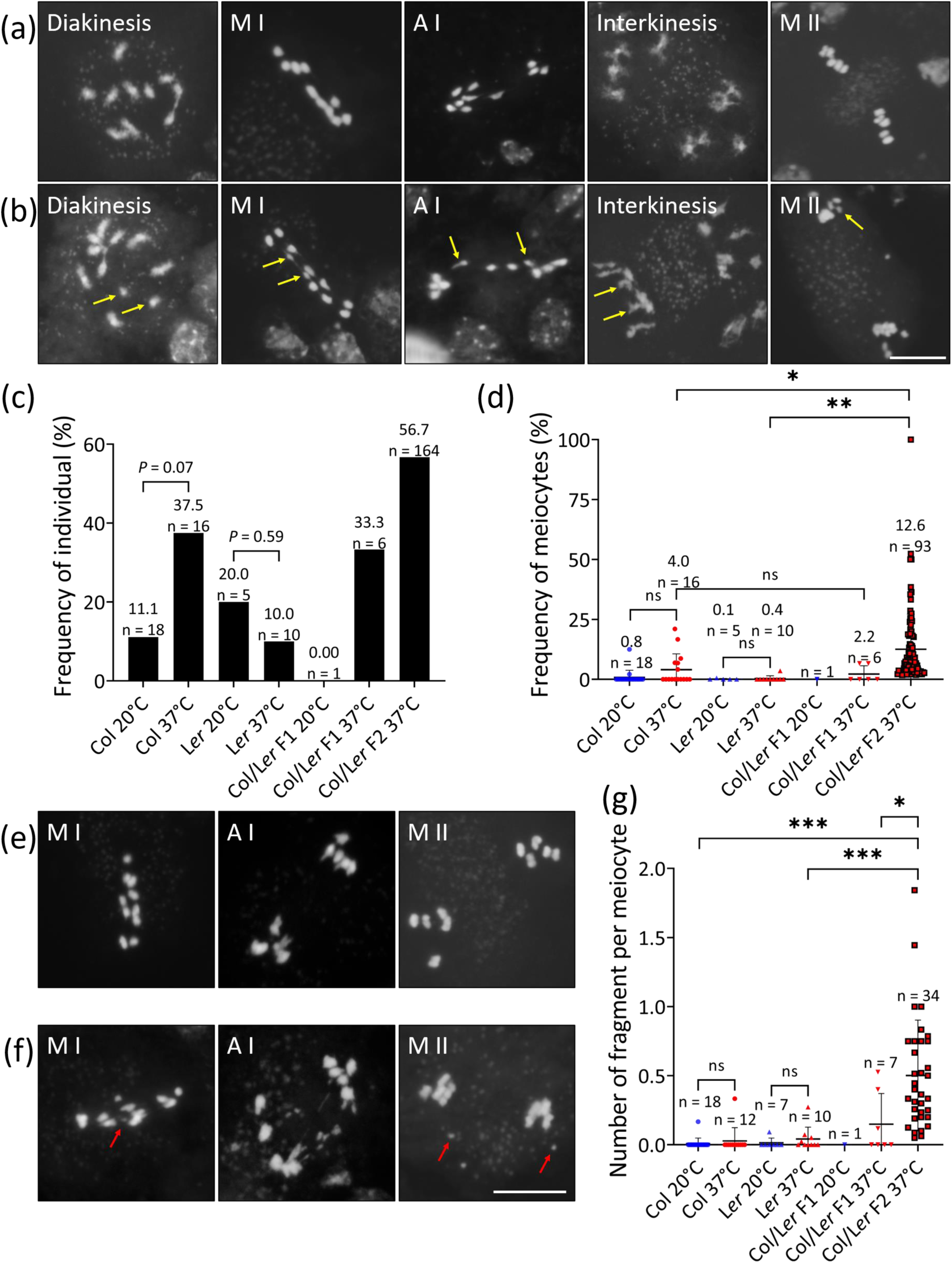
The Col/L*er* F2 population shows a higher level of chromosome instability under heat stress. (a) and (b) Representative images of diakinesis, metaphase I, anaphase I, interkinesis and metaphase II meiocytes showing normal (a) or pre-mature segregation of sister chromatids (b) under heat stress. The yellow arrows indicate separated sister chromatids. (c) Graph showing the rates of individuals that show abnormal segregation of sister chromatids in Col, L*er*, Col/L*er* hybrids and Col/L*er* F2 at 20°C or 37°C. The indicated significance levels were determined based on χ^2^-tests. (d) Graph showing the rates of meiocytes exhibiting premature separation of sister chromatids in Col, L*er*, Col/L*er* hybrids and Col/L*er* F2 at 20°C or 37°C. (e and f) Representative images of metaphase I, anaphase I and metaphase II chromosomes with normal (e) or interfered integrity (f) under heat stress. The red arrows indicate chromosome fragments. (g) Graph showing the average numbers of chromosome fragment per meiocyte in Col, L*er*, Col/L*er* hybrids and Col/L*er* F2 at 20°C or 37°C. Stars indicate significance levels based on unpaired *t* tests; frequencies indicate the rates of the corresponding phenotypes; n indicates the number of individuals; ***, *P* < 0.001; **, *P* < 0.01; *, *P* < 0.05; ns, *P* > 0.05; scale bars, 10 μm.

### Homolog pairing and synapsis facilitate timely sister-chromatids separation under heat stress

We then related homolog pairing and bivalent formation with the phenotype of premature sister-chromatids separation in our heat-stressed Col/L*er* F2 population. A positive but weak correlation was detected between the fraction of meiocytes showing cohesion defect with the fraction of meiocytes showing partial pairing, or the diakinesis meiocytes producing an average of one bivalent (Fig 8a and b). Notably, among the meiocytes showing ectopic sister-chromatid separation, the meiocytes in metaphase I, anaphase I or metaphase II stages occupied 8.7%, 35.8% and 59.0%, respectively (Fig 8c). This implied that heat stress primarily destabilizes centromeric cohesion in late or after meiosis I. Moreover, we found a strong and positive correlation between the fraction of meiocytes exhibiting chromosome fragmentations with meiocytes in pachytene undergoing normal homolog pairing (Fig 8d) and with meiocytes in diakinesis producing five bivalents (Fig 8e).

**Figure 8.**
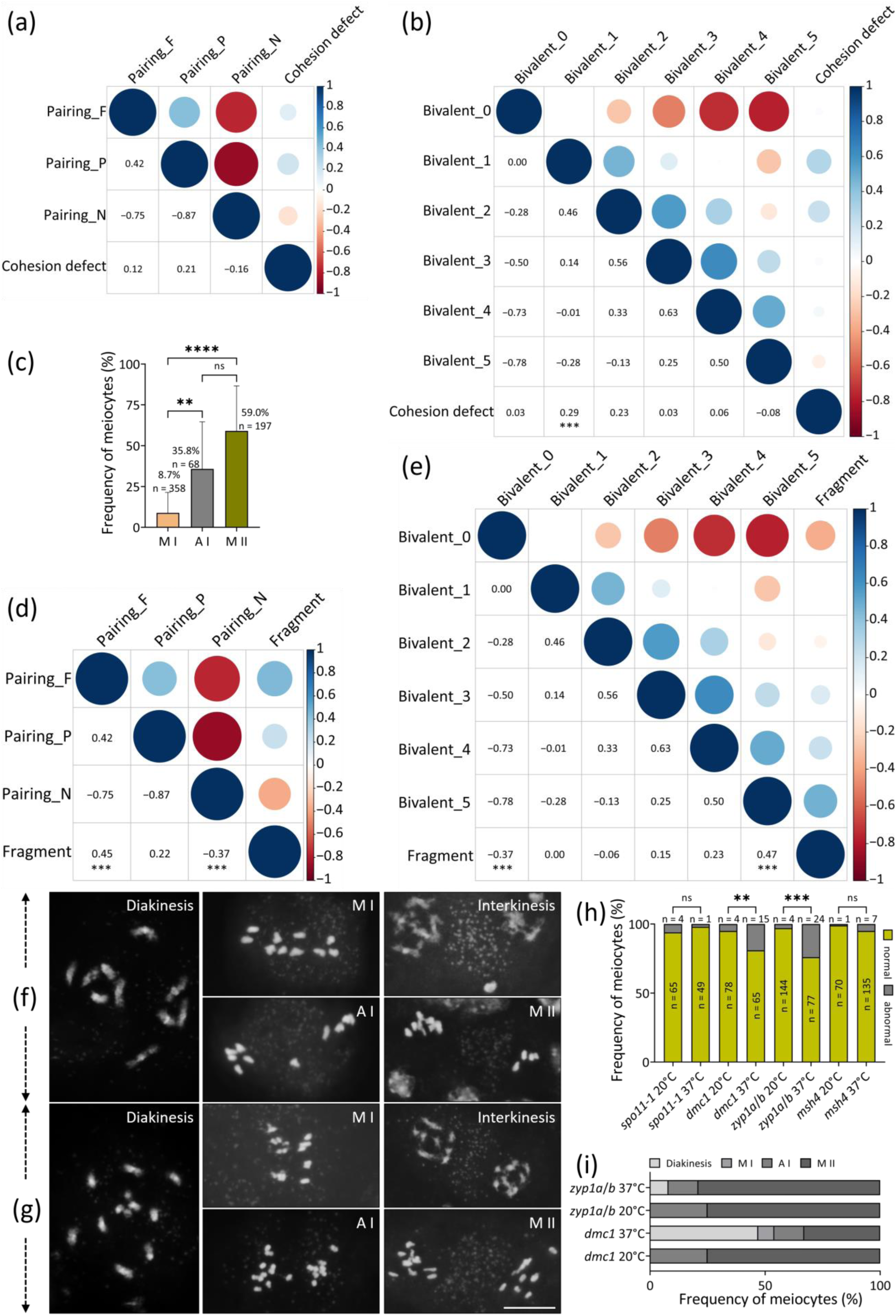
Homolog pairing and synapsis facilitate timely sister-chromatids separation under heat stress. (a and b) Graphs showing the correlation between the rates of meiocytes showing different pairing status (a) or bivalent formation (b) with the frequency of meiocytes with cohesion defect in the Col/L*er* F2 population under heat stress. (c) Graph showing the rates of metaphase I, anaphase I and metaphase II meiocytes with cohesion defect in the Col/L*er* F2 population under heat stress. (d and e) Graphs showing the correlation between the frequencies of meiocytes showing different pairing status (d) or bivalent formation (e) with the rate of meiocytes showing chromosome fragmentation in the Col/L*er* F2 population under heat stress. Spearman correlation analysis was performed. Numbers indicate the correlation coefficients between the corresponding phenotypes; significance of correlation was calculated using *t*-tests; the significance levels in the cells without indication of significance values were not calculated. (f and g) Representative images of diakinesis, metaphase I, anaphase I, interkinesis and metaphase II meiocytes showing normal (f) or untimely sister-chromatids separation (g). (h) Graph showing the rates of meiocytes with normal or untimely sister-chromatids separation in *spo11-1*, *dmc1*, *zyp1a*/*b* and *msh4* at 20°C and 37°C. (i) Graph showing the rates of different meiosis-staged meiocytes exhibiting untimely sister-chromatids separation in *dmc1* and *zyp1a*/*b* at 20°C and 37°C. For (c) and (h), χ^2^-tests were performed; frequencies indicate the rates of the corresponding phenotypes; n indicates the number of analyzed cells; ***, *P* < 0.001; **, *P* < 0.01; ns, *P* > 0.05; scale bar, 10 μm.

To further investigate the impact of homolog pairing and/or synapsis and CO formation on sister-chromatids cohesion under heat stress, we analyzed a *spo11-1* mutant (Col background), in which DSB formation is largely abolished and homolog pairing and recombination are impaired (56). Untimely separation of sister chromatids was observed in *spo11-1* at both temperatures, but there was no significant difference (Fig 8f-h, *P* > 0.05). Next, we analyzed the *dmc1* (Col background), in which DSBs are formed but homolog pairing and CO formation do not occur (57), and the double *zyp1a*/*b* mutant (Col background), in which homolog pairing occurs but synapsis is impaired (14, 58, 59). Both *dmc1* and *zyp1a*/*b* exhibited premature sister-chromatids segregation at 20°C, and, the fraction of meiocytes showing the cohesion defect in both mutants were significantly increased under heat stress (Fig 8f-h; *P* < 0.01 for *dmc1* and *P* < 0.001 for *zyp1a*/*b*). Interestingly, the defect-harboring meiocytes in *dmc1* occurred by 66.7% at and/or prior to anaphase I, while, in *zyp1a*/*b*, the irregularity was found by 79.2% at metaphase II (Fig 8i). These findings implied that DMC1- and ZYP1-dependent homolog pairing and/or synapsis play a role in facilitating timely separation of sister chromatids under heat stress, yet possibly via different mechanisms. To address if the distinct distribution of differently-staged meiocytes showing ectopic sister-chromatids separation between *dmc1* and *zyp1a*/*b* was owing to the difference in CO formation, we examined a *msh4* mutant (Col background), which undergoes normal homolog synapsis but loses approximately 85% COs (60). No significant difference in the rates of meiocytes with cohesion defect was detected at 20°C and 37°C (Fig 8h, *P* > 0.05).

### Chromosome integrity in both Col and L*er* under heat stress relies on functional ATM

We previously reported that ATM-controlled DSB repair is required for maintenance of meiotic chromosome integrity in Col under heat stress (51). Thus, we decided to examine whether the role of ATM in protecting chromosome stability under heat stress varies between Col and L*er*. To this end, we took advantage of an *atm* mutant allele, *atm-5*, which is in L*er* background and shows defects in DSB repair (61). We found that, as the case in *atm-2*, an allele in Col background (51), chromosome fragments in *atm-5* was significantly increased after heat treatment (Fig 9a and b; *P* < 0.001). A bi-allelic *atm* mutant *ATM^atm-2^*^/*atm-5*^, which harbored the *atm-2* and *atm-5* alleles on either homolog, was generated, and behaved similarly as *atm-2* and *atm-5* under heat stress (Fig 9c and d). We also examined the impact of heat on homolog pairing in *atm-5*. At 20°C, most pachytene meiocytes showed normal pairing and synapsis of homologs, and 16.9% of meiocytes showed a partial pairing phenotype (Fig 9e and g). At 37°C, most pachytene meiocytes in *atm-5* showed a partial pairing phenotype, with only 3.9% cells showing an impaired homolog pairing (Fig 9e and g), similar to the case in heat-stressed L*er* (Fig 3j). Immunolocalization revealed an increased level of patchy ZYP1 configuration in *atm-5* compared with that in L*er* (Fig 9f and h; Fig 3t), which was likely caused by the disrupted chromosome integrity due to ATM dysfunction (51). Similar alterations in homolog pairing and SC formation were observed in *ATM^atm-2^*^/*atm-5*^ at either temperature (Fig 9i-l). These data suggest that there is no accession-specific difference in dependency of ATM-mediated DSB repair for meiotic chromosome integrity under heat stress.

**Figure 9.**
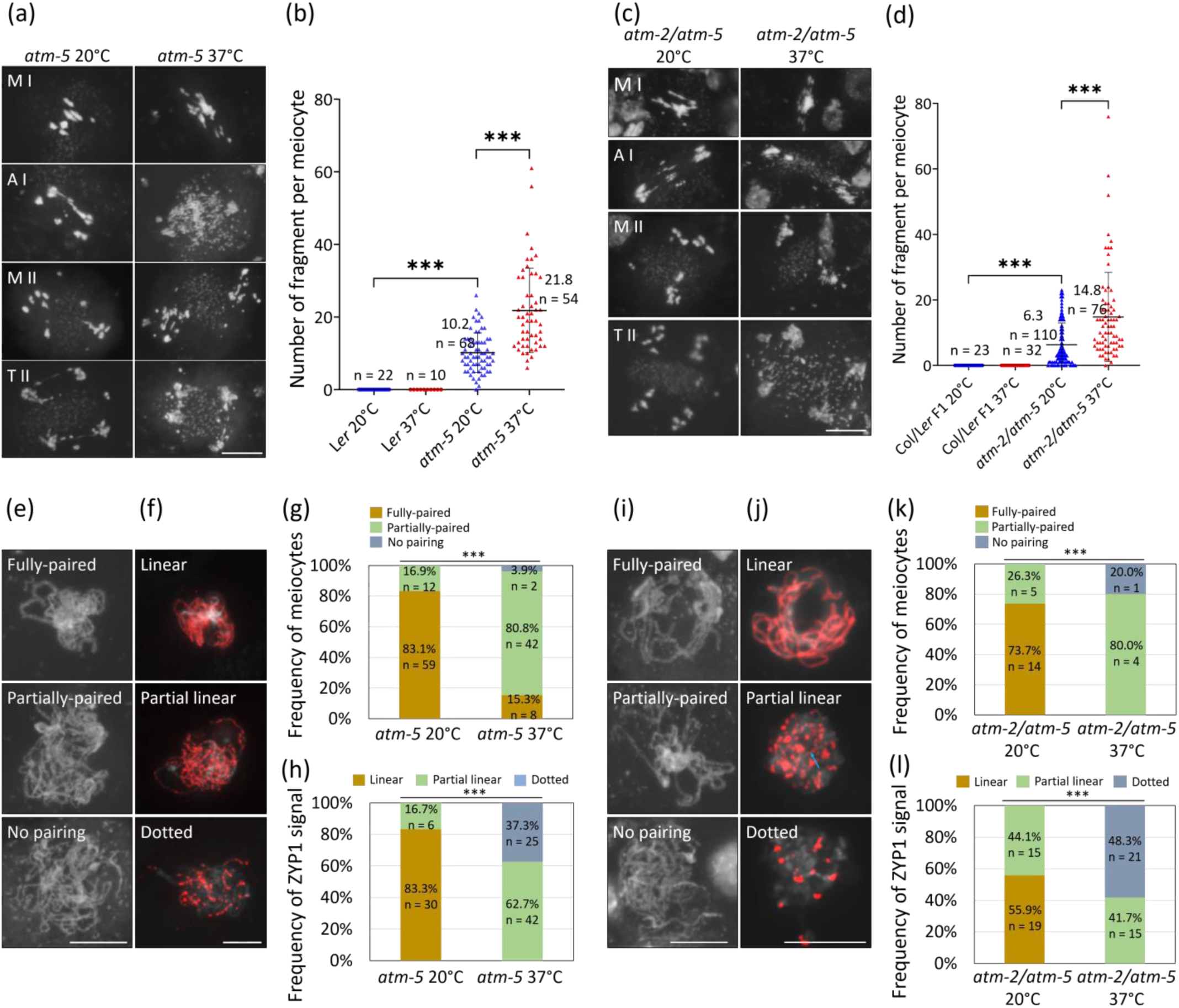
Chromosome integrity under heat stress in both Col and L*er* relies on ATM. (a and c), Metaphase I, anaphase I, metaphase II and telophase II chromosomes in *atm-5* (a) and *ATM^atm-2^*^/*atm-5*^ (c) at 20°C and 37°C. (b and d), Graphs showing the average number of chromosome fragment in *atm-5* (b) and *ATM^atm-2^*^/*atm-5*^ (d) at 20°C and 37°C. Significance levels are shown based on unpaired *t* tests. (e and i), Representative images of pachytene chromosomes showing fully-paired, partially-paired, or no pairing in *atm-5* (e) and *ATM^atm-2^*^/*atm-5*^ (i) at 20°C and 37°C. (f and j), Representative images of immunolocalization of ZYP1 on pachytene chromosomes showing linear, partial linear, or patchy phenotypes in *atm-5* (f) and *ATM^atm-2^*^/*atm-5*^ (j) at 20°C and 37°C. The blue arrow indicates linear ZYP1 signals. (g and k), Graphs showing the rates of different pairing status of pachytene chromosomes in *atm-5* (g) and *ATM^atm-2^*^/*atm-5*^ (k) at 20°C and 37°C. Significance levels are shown based on χ^2^-tests. (h and l), Graphs showing the rates of different ZYP1 configurations on pachytene chromosomes in *atm-5* (h) and *ATM^atm-2^*^/*atm-5*^ (l) at 20°C and 37°C. Significance levels are shown based on χ^2^-tests; frequencies or numbers indicate the rate or quantity of the corresponding phenotypes; n indicates the number of analyzed cells; ***, *P* < 0.001; scale bars, 10 μm.

## Discussion

Our previous study demonstrated that heat stress reduces DSB formation and disrupts pairing as well as synapsis of homologs in Col (36, 37). In this study, we report that Col and L*er* have a striking difference in heat tolerance of meiotic recombination. Specifically, heat stress does not impair SC formation and does not reduce the numbers of RAD51 or DMC1 foci in L*er* (Fig 2 and 3). Based on our previous studies, the distinct heat tolerance of DSB formation between Col and L*er* is likely due to a difference in the activity and/or localization of the recombination proteins, and not in transcription levels (36, 51). Since the number of bivalents is strongly correlated with homolog pairing status (Fig 5), it is likely that heat suppresses meiotic recombination predominantly by interfering with DSB formation and/or the RAD51- and DMC1-mediated homolog pairing. In support, DMC1 has been found to play a role in maintaining CO formation at high temperature in wheat (62, 63). Nevertheless, since there is still a low level of DSBs in heat-stressed Col (Fig 2), and some meiocytes in L*er* display compromised homolog pairing under heat stress (Fig 3), we propose that heat can also directly attenuate homolog pairing apart from reducing DSB formation (36, 37, 51, 64). Based on the analysis of Col/L*er* hybrids, we conclude that the heat tolerance of meiotic recombination in L*er* is a dominant trait (Fig 4). 10% ∼ 20% of F2 offspring behaving as L*er* suggests that the dominance relies on the presence of several alleles. While all alleles are present in Col/L*er* hybrids, the linkage of these alleles is broken in the meiosis of these hybrids leading to a differentiated pattern of heat tolerance in the F2 population (Fig 5).

Heat induces an increased fraction of interkinesis meiocytes displaying a non-condensed chromosome configuration in Col, which does not occur in L*er* and the Col/L*er* F1 hybrids, but occurs again, and, at the same level, in the F2 population (Fig 6d). Meanwhile, heat increases the rate of interkinesis meiocytes exhibiting a less-condensed phenotype in some F2 individuals but not in inbreds or hybrids, which suggests a higher heat sensitivity in F2 offspring (Fig 6e). The heat-induced abnormalities in interkinesis phenotypically mimic slowed meiosis progression or a failure of entering into meiosis II, which implies that meiosis progression and/or cell cycle transition may be perturbed possibly due to alterations in expression and/or function of *TAM/CYCA1;2* (*TAM*) or *OMISSION OF SECOND DIVISION 1* (*OSD1*), or other meiotic cell cycle regulators (36, 65–67). Furthermore, correlation analysis suggests that meiotic recombination or its related regulators may impact meiotic cell cycle progression and/or chromosome morphology after anaphase I beyond its impact on prophase I progression under heat stress (Fig 6) (25).

Incorrect sister-chromatids separation in the F2 offspring (Fig 7) implies that heat perturbs the function of the cohesion complex. It has recently been reported that high temperatures reduce the amounts of CENH3 in Arabidopsis (35), which may lead to interfered centromere function and cohesion instability. Notably, we found that most meiocytes showing cohesion defects are in anaphase I or metaphase II (Fig 8), suggesting that heat primarily perturbs centromeric cohesion during anaphase I and interkinesis. In Arabidopsis, SHUGOSHIN 1 (SGO1) and PROTEIN PHOSPHATASE 2A (PP2A), and the Arabidopsis homolog of securin PATRONUS (PANS1) protect meiotic centromeric cohesion at anaphase I and diakinesis, respectively (68–71). In addition, PP2A is co-expressed with genes involved in response to abiotic stresses including heat in multiple developmental stages in Arabidopsis (72). It is thus possible that heat stress alters the expression and/or function of these cohesion regulators. In multiple species, bivalent formation is required for correct orientation of sister kinetochores and thus balanced chromosome segregation in meiosis I (73–78). Our correlation analysis suggests that homolog pairing and/or synapsis but possibly not chiasmata formation may directly or indirectly impact timely sister-chromatids separation under heat stress (Fig 8). However, heat-stressed *spo11-1* does not show a higher level of defective sister-chromatids separation (Fig 8), suggesting that the effect of heat stress on homolog pairing-mediated cohesion stability may rely on DSB formation, and, the increased aneuploid nuclei due to chromosome mis-segregation observed in heat-stressed *spo11-2* are not owing to premature sister-chromatids separation (35).

We observed an increase of chromosome fragments in some Col/L*er* F2 individuals but not in either inbreds or hybrids under heat stress (Fig 7), suggesting that DSB repair is perturbed in those fragment-harboring F2 plants. This phenotype, together with the increased fraction of meiocytes showing premature sister-chromatid separation, imply that transgressive effects become visible in the F2 population showing defects exceeding that observed in parents. ATM-mediated DSB repair is required for meiotic chromosome integrity in both Col and L*er* under heat stress (Fig 9) (51), dysfunction of which might be one of the mechanisms that underpin the heat-interfered chromosome integrity in those F2 individuals. The upstream factor and/or signaling pathway that regulates ATM expression or activity in DSB repair in response to heat stress require further investigation. Our data suggest that there is no accession-specific difference in the dependency of ATM-mediated DSB repair for maintenance of meiotic chromosome integrity under heat stress. Alternatively, ATM-mediated mechanism acts in an epistatic manner to protect meiotic chromosome stability under heat stress. Interestingly, the fragmentation phenotype is strongly and positively correlated with successful homolog pairing and bivalent formation (Fig 8d and e). We speculate that these associations are possibly a subsequent effect of heat stress on DSB formation and repair, the heat tolerance of which are regulated via different pathways. Specifically, in the individuals that show normal homolog pairing and CO formation, DSB formation likely occurs normally, however, DSB repair may be perturbed, which consequently leads to lesions in chromosome integrity. On the contrary, the individuals with impaired homolog pairing and CO formation likely have a significantly reduced DSB formation, which compensates for the perturbation of DSB repair, and thus show relatively undisturbed chromosome integrity. Thus, DSB formation and homolog pairing likely are the predominant targets, which facilitate meiosis progression and genome stability at post-recombination stages under heat stress. Taken together, our study reveals that natural variation in multiple genomic loci controls heat tolerance of meiosis in Arabidopsis, holding a great potential to be exploited in breeding programs.

## Material and Methods

### Plant materials and growth conditions

*Arabidopsis thaliana* (L.) accessions Columbia-0 (Col-0) and Landsberg *erecta* (L*er*), the *atm-2* (50), *atm-5* (61), *er-105*, *mpk6*, *er-105 mpk6* (52), *dmc1* (79), *spo11-1-3* (56), *msh4* (60) and *zyp1a*/*b* (14) mutants were used in this study. For simplification ‘Col’ was used throughout the main text. Col/L*er* hybrids were obtained by crossing using L*er* as pollen donator. The bi-allelic mutant *ATM^atm-2^*^/*atm-5*^ was obtained by crossing the heterozygous *atm-2* and *atm-5* mutants. Genotyping was performed using the primers listed in Table S1. Arabidopsis were cultivated in a growth chamber with a 16 h day/8 h night and 20°C condition. For heat treatment, young flowering plants were treated by 37°C for 24 h in a humid chamber with a 16 h day/8 h night condition. To avoid potential impact of circadian clock, heat treatment was applied starting from 8:00-10:00 am.

### Cytological analysis of meiocytes

After heat treatment, plants were transferred back to 20°C for 6-12 h recovery, and tetrad stage meiocytes were analyzed (38). A drop of 4.5% (w/v) lactopropionic orcein staining buffer was added to the surface of a tetrad-staged flower bud (floral bud stage 9, 0.3-0.4 mm) (80) followed by squashing to release meiocytes.

### Preparation of chromosome spreads

Inflorescences were isolated and fixed in cold Carnoy’s fixative for at least 24 h. Flower buds at stages beyond tetrads were removed, and the remaining buds were washed twice in distilled water and once in citrate buffer (10 mM, pH = 4.5), followed by incubation in a digestion enzyme mixture (0.3% pectolyase and 0.3% cellulase in citrate buffer) at 37°C for 3 h. Digested flower buds were subsequently washed once in distilled water, which thereafter were macerated in distilled water on a glass slide. Two rounds of 60% acetic acid were added to the slide, which was dried on a hotplate at 45°C. The slide was flooded with cold Carnoy’s fixative and then was air dried. 4’,6-diamidino-2-phenylindole (DAPI) was diluted to 5 μg/mL in Vectashield antifade mounting media (Vector Laboratories).

### Immunolocalization of ɑ-tubulin and meiotic recombination proteins

To perform immunolocalization of ɑ-tubulin, inflorescences were fixed by 4% (w/v) paraformaldehyde under vacuum for 30 min, followed by twice washes in distilled water and once in citrate buffer (10 mM, pH = 6.0). Meiosis-staged flower buds were digested in an enzyme mixture consisting of 0.3% (w/v) cellulase (Sigma), 0.3% (w/v) pectolyase (Sigma) and 0.3% (w/v) cytohelicase (Sigma) in a humid chamber at 37°C for 3 h. After the digestion, anthers were dissected, squashed, and fixed on a slide by freezing in liquid nitrogen. Released cells on the slide were immobilized with a thin layer of gel containing 1% (w/v) gelatin, 1% (w/v) agarose and 2.5% (w/v) sucrose. After rinsing with phosphate buffered saline with 1% (v/v) Triton-X 100 for 30 min, the ɑ-tubulin antibody diluted in phosphate buffered saline with 0.1% (v/v) Triton X-100 and 4.5 g/L bovine serum albumin was added and the slide was incubated overnight at 4°C. The slide washed three times with phosphate buffered saline with 0.02% tween-20 was incubated with a secondary antibody at 37°C for 1 h in the dark. Immunolocalization of meiotic recombination proteins was performed as previously described (81, 82), and the antibodies used in this study have been reported previously (36, 38, 51).

### Quantification of fluorescent foci, fluorescence intensity and chromosome fragments

Fluorescent images were taken by an Olympus IX83 inverted fluorescence microscope with a X-Cite lamp and a Prime BSI camera, and Z-stacks were processed using ImageJ. The fluorescent foci merged onto DAPI-stained chromosomes were counted. For the pixel intensity plot, the pixel brightness through a region of interest was measured using ImageJ and plotted against the X dimension. To score the number of chromosome fragments, chromosome bodies in meiocytes at metaphase I, anaphase I and metaphase II that were obviously brighter than organelles were counted followed by a subtraction of expected chromosome number (5 at metaphase I, 10 at anaphase I or metaphase II).

### Statistical analysis

Significance was calculated using unpaired *t*-tests (Fig 1b; Fig 2c, d, h and j; Fig 4b, d, and h; Fig 6d and e; Fig 7d and g; Fig 9b and d) or Chi-squared tests (Fig 1f; Fig 2f; Fig 3j, l and t; Fig 4f, j, l and n; Fig 7c; Fig 8c and h; Fig 9g, h, k and l) with the software GraphPad Prism. The significance level was set as *P* < 0.05. To calculate the frequency of individuals or meiocytes showing either altered chromosome morphology in interkinesis (Fig 6c, d and e), untimely sister-chromatids segregation, or number of chromosome fragments (Fig 7c, d and g), the individuals, in which a meiocyte with the corresponding defect was observed, and at least three meiocytes at the corresponding stage were found, were used for calculation. The numbers of analyzed meiocytes or individuals are shown in the figures or main text. Correlation between different meiosis phenotypes was determined based on Spearman’s rank correlation tests. Coefficients were calculated using Corrplot package on the R statistical platform, and Ggpubr package was used for visualization.

## Acknowledgement

The authors thank the National Natural Science Foundation of China (32000245), Hubei Provincial Natural Science Foundation of China (2024AFB695), the Fundamental Research Funds for the Central Universities of South-Central Minzu University (CZZ24011), Knowledge Innovation Program of Wuhan-Shuguang Project (2022020801020410) and general funds of University of Hamburg (to A.S.) for the financial support on this study. They thank Pingli Lu (Henan University) for sharing the *atm-5* mutants. They thank Chao Yang (HZAU) for providing the *zyp1a*/*b* mutant. They thank Yuan Qin (FAFU) for sharing the *er-105*, *mpk6* and the *er-105 mpk6* mutants.

## Author contributions

**Conceptualization:** Bing Liu.

**Data curation:** Jiayi Zhao, Bing Liu.

**Formal analysis:** Jiayi Zhao, Zhengze Wang, Ziming Ren, Zhihua Wu and Xiaoning Lei.

**Funding acquisition:** Arp Schnittger, Bing Liu.

**Investigation:** Jiayi Zhao, Huiqi Fu, Min Zhang, Yaoqiong Liang, Xueying Cui, Wenjing Pan, Yujie Zhang, Xin Gui, Li Huo, Chong Wang.

**Methodology:** Jiayi Zhao, Arp Schnittger, Wojciech P. Pawlowski, Bing Liu.

**Project administration:** Bing Liu.

**Resources:** Bing Liu.

**Supervision:** Bing Liu.

**Validation:** Jiayi Zhao, Bing Liu.

**Writing - original draft:** Bing Liu.

**Writing - review & editing:** Jiayi Zhao, Huiqi Fu, Arp Schnittger, Wojciech P. Pawlowski, Bing Liu.

## Data availability

All data supporting the conclusions drawn in our study are fully documented in this article and Supporting Information

(including Fig S1–S4; Table S1).

## Accession numbers

Accession numbers of genes studied in this work are: *SPO11-1* (AT3G13170), *RAD51* (AT5G20850), *DMC1* (AT3G22880), *ASY4* (AT2G33793), *ZYP1A* (AT1G22260), *ZYP1B* (AT1G22275), *HEI10* (AT1G53490), *ERECTA* (AT2G26330), *MPK6* (AT2G43790), *ATM* (AT3G48190) and *MSH4* (AT4G17380).

## Supporting Information

Fig S1. Heat stress does not affect microtubule organization during prophase I in *Arabidopsis thaliana*.

Fig S2. Flower architectures in Col, L*er* and Col/L*er* hybrids.

Fig S3. Correlation analysis of synapsis, bivalent formation, and segregation of homologs at anaphase I in Col, L*er* and Col/L*er* hybrids.

Fig S4. Representative images of meiotic chromosomes from anaphase I to metaphase II in wild-type Arabidopsis.

Table S1. Primers used in this study.

## Notes

### Competing Interest Statement

The authors have declared no competing interest.

### Summary of Updates

an author summary has been included in the revised version

