## Supporting Information for "Genetic variation in the species *Arabidopsis thaliana* reveals the existence of natural heat resilience factors for meiosis"

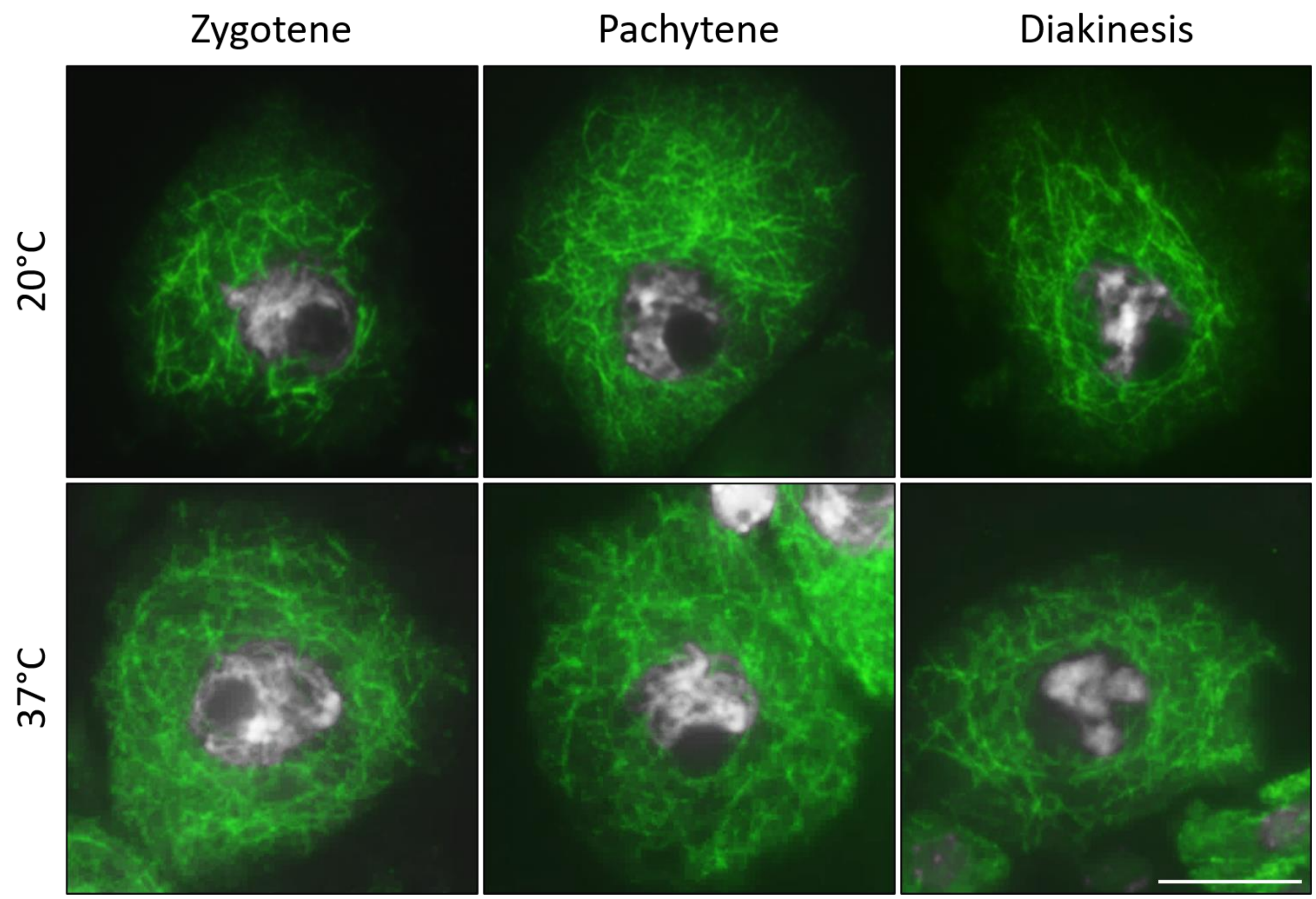


Fig S1. Heat stress does not affect microtubule organization during prophase I in *Arabidopsis thaliana*. White, DAPI; green, α-tubulin. Scale bar = 10 μm.

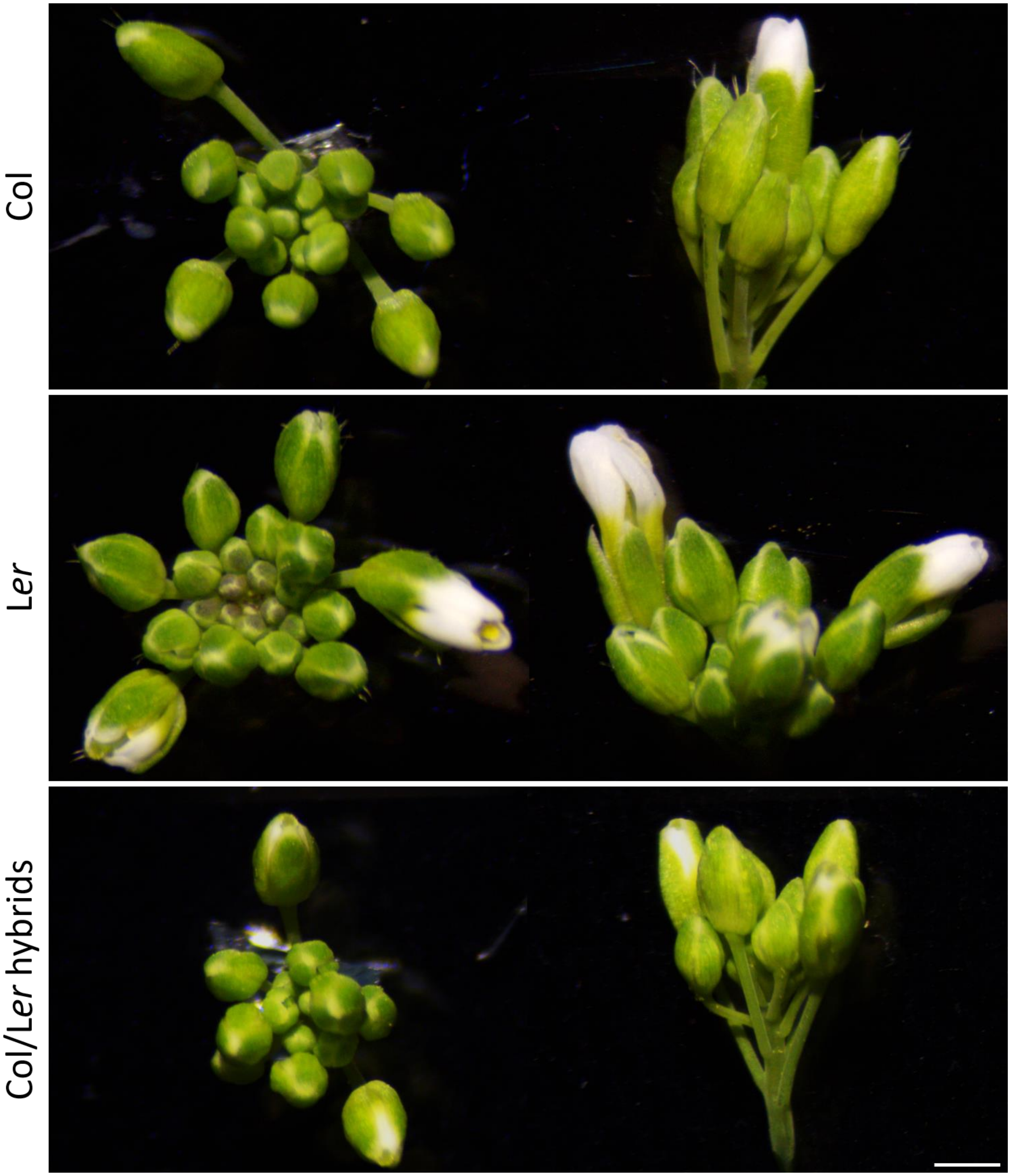


Fig S2. Flower architectures in Col, L*er* and Col/L*er* hybrids. Scale bar = 1 mm.

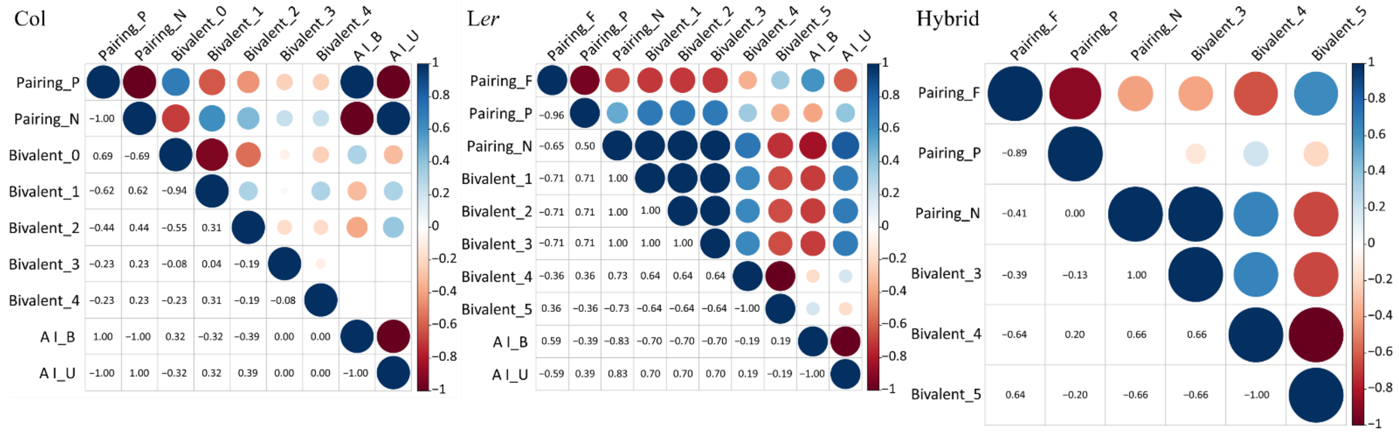


Fig S3. Correlation analysis of synapsis, bivalent formation, and segregation of homologs at anaphase I in Col, L*er* and Col/L*er* hybrids. Spearman correlation analysis was performed. Numbers indicate the correlation coefficients between the corresponding phenotypes; unindicated significances in the cells mean that the significances were not calculated.

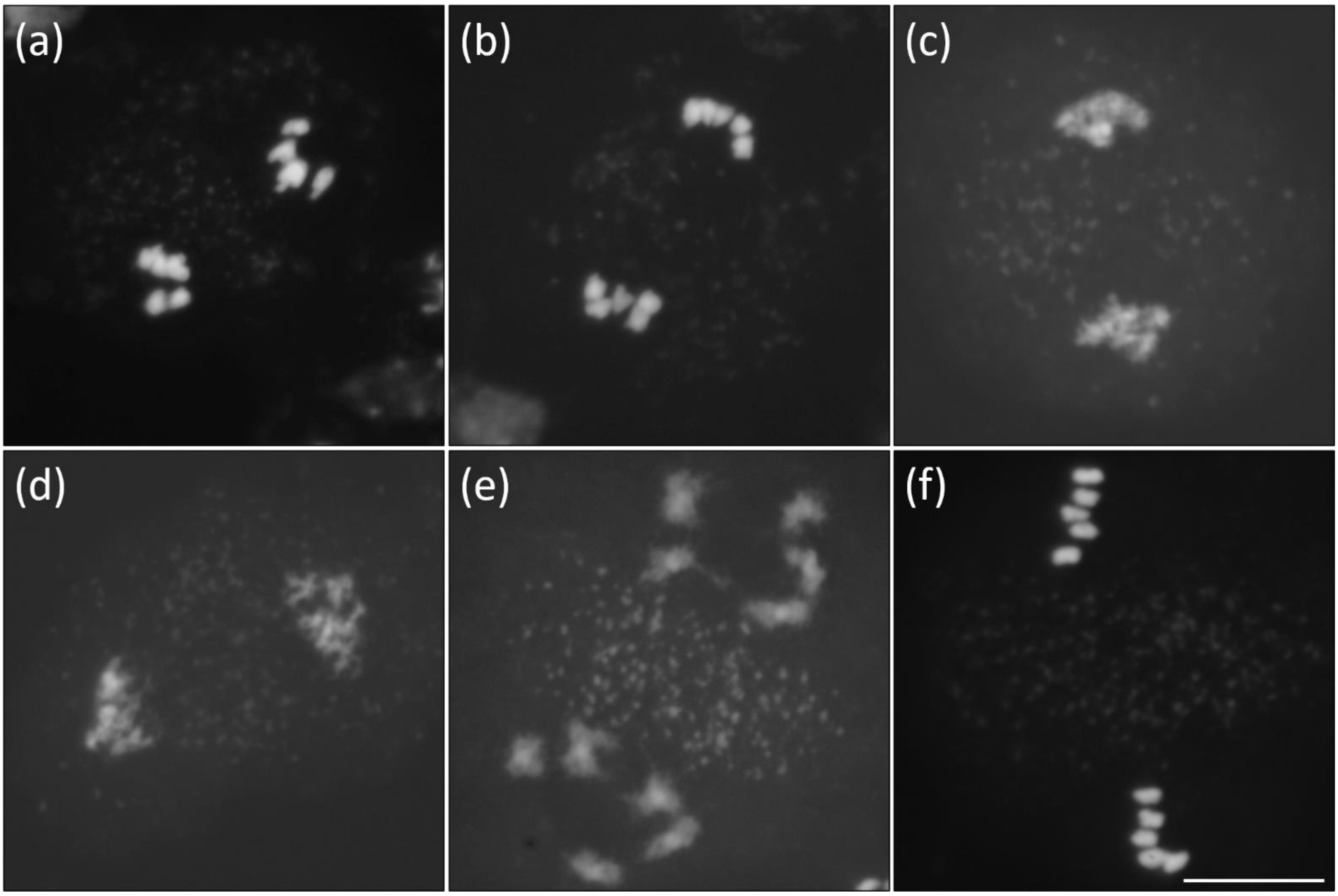


Fig S4. Representative images of meiotic chromosomes from anaphase I to metaphase II in wild-type Arabidopsis. (a-f) Middle anaphase I showing segregating homologs (a), mature anaphase I (b), early interkinesis showing decondensed (c) and recondensing homologs (d), mature interkinesis (e) and metaphase II (f) meiocytes in wild-type *Arabidopsis thaliana*.

Table S1. Primers used in this study.

| Primers | Sequence (5' - 3') | Purpose |
|---|---|---|
| atm-2 F | ATCCATGTGGTTCAGTCTTGC | *atm-2* genotyping |
| atm-2 R | TTGGTATCCTGCAGAGGAAAG | |
| atm-5 F | GTTGCCGGATCAGCAGTAGT | *atm-5* genotyping |
| atm-5 R | GTGCCCTAGTCCAGTTTCCC | |
| er-105 F | AAGAAGTCATCTAAAGATGTGA | *er-105* genotyping |
| er-105 R | AGCTGACTATACCCGATACTGA | |
| mpk6 F | CTCTGGCTCATCGCTTATGTC | *mpk6* genotyping |
| mpk6 R | ATCTATGTTGGCGTTTGCAAC | |
| msh4 F | CGGCTTCACTGCATCTATCTC | *msh4* genotyping |
| msh4 R | TGGAATGGATCAATGAGTTCC | |
| zyp1a F | GCAATGAAAACGAGAAGCAGT | *zyp1a* genotyping |
| zyp1a R | TACCTGGTCGTTCCTCTGTG | |
| zyp1b F | CTCGCATTTGCTGGTTTAAAGAGTC | *zyp1b* genotyping |
| zyp1b R | TGCGTATATTGCTAGGTTTATATTG | |
| dmc1 F | CCTGCAATGGTCTCATGATGCATAC | *dmc1* genotyping |
| dmc1 R | GATGCAATCGATATCAGCCAATTTTAGAC | |
| C/L Chr 1-1 F | CTCTTGGTGGTGTCCCAAGT | Genotyping chr 1 in Col/L*er* F1 and F2 |
| C/L Chr 1-1 R | TCGACGCAGTTTTTCATCAG | |
| C/L Chr 1-2 F | ACAAAATGCCGATCCAACAT | |
| C/L Chr 1-2 R | TGCTGAAAACGTCAAGACCA | |
| C/L Chr 2-1 F | GTTTGGATCAGTCCCAGCTC | Genotyping chr 2 in Col/L*er* F1 and F2 |
| C/L Chr 2-1 R | TGAAAAAGTGGTGGAACCAA | |
| C/L Chr 2-2 F | TTGGGTTTGAGTCACATTCG | |
| C/L Chr 2-2 R | TACCTCCAACAAGCCACACA | |
| C/L Chr 3-1 F | TTCAGCAACCTTCGATAAATCA | Genotyping chr 3 in Col/L*er* F1 and F2 |
| C/L Chr 3-1 R | CCATTGCCACCGTAGAAACT | |
| C/L Chr 3-2 F | GGTTTGGTGGGAGAGAATGA | |

| | | |
|---|---|---|
| C/L Chr 3-2 R | CAAAAGAAATGCAACGAGACA | |
| C/L Chr 4-1 F | TTTTTAATTAAGGGAACAAAATGGA | |
| C/L Chr 4-1 R | TTGTGTCATATGTCAAGTCTGTCG | |
| C/L Chr 4-2 F | CGGCGACTGTGAATTATGTG | Genotyping chr 4 in Col/L*er* F1 and F2 |
| C/L Chr 4-2 R | CCGTCACAATCCTGACTCAA | |
| C/L Chr 5-1 F | AGCTCAAGCAATCCAACCAC | |
| C/L Chr 5-1 R | TCCAGCTTTGGACTTCTTCG | Genotyping chr 5 in Col/L*er* F1 and F2 |
| C/L Chr 5-2 F | CCCAATACCGAATCAATCAAA | |
| C/L Chr 5-2 R | GGGAACTGATTGGCTGACAC | |
